# The stretch-shortening cycle effect is not associated with cortical or spinal excitability modulations

**DOI:** 10.1101/2024.08.19.608542

**Authors:** Lea-Fedia Rissmann, Brent James Raiteri, Wolfgang Seiberl, Tobias Siebert, Daniel Hahn

## Abstract

It is unclear whether cortical and spinal excitability modulations contribute to enhanced SSC performance. Therefore, this study investigated cortical and spinal excitability modulations during and following shortening of stretch-shortening cycle (SSC) contractions compared with pure shortening (SHO) contractions. Participants (N = 18) performed submaximal voluntary plantar flexion contractions while prone on the dynamometer bench. The right foot was strapped onto the dynamometer’s footplate attachment and the resultant ankle joint torque and crank arm angle were recorded. Cortical and spinal excitability modulations of the soleus muscle were analyzed by eliciting compound muscle actional potentials via electrical nerve stimulation, cervicomedullary motor-evoked potentials (CMEPs) via electrical stimulation of the spinal cord, and motor-evoked potentials (MEPs) via magnetic stimulation of the motor cortex. Mean torque following stretch was significantly increased by 7±3% (*p*=0.029) compared with the fixed-end reference (REF) contraction and mean torque during shortening of SSC compared with SHO was significantly increased by 12±24% (*p*=0.046). Mean steady-state torque was significantly lower by 13±3% (*p*=0.006) and 9±12% (*p*=0.011) following SSC compared with REF and SHO, respectively. Mean steady-state torque was not significantly lower following SHO compared with REF (7±8%, p=0.456). CMEPs and MEPs were also not significantly different during shortening of SSC compared with SHO (*p*≥0.885) or during the steady state of SSC, SHO, and REF (*p*≥0.727). Therefore, our results indicate that SSC performance was not associated with cortical or spinal excitability modulations during or after shortening, but rather driven by mechanical mechanisms triggered during active stretch.

**Key points:**

1. A stretch-shortening cycle (SSC) effect of 12% was observed during EMG-matched submaximal voluntary contractions of the human plantar flexors
2. The SSC effect was neither associated with cortical or spinal excitability modulations nor with stretch-reflex activity
3. The SSC effect was likely driven by mechanical mechanisms related to active muscle stretch, which have long-lasting effects during shortening
4. Residual force depression following SSC was not attenuated by the long-lasting mechanical mechanisms triggered during active muscle stretch
5. Steady-state torques were lower following shortening of SSCs versus pure shortening and fixed-end contractions at the same final ankle joint angle, but the torque differences were not correlated with cortical or spinal excitability modulations

## Introduction

During everyday locomotion, lower limb muscle-tendon units (MTU) often stretch before shortening, which is known as a stretch-shortening cycle (SSC). SSCs are unique because torque, work and power outputs during and after the shortening phase are typically higher than shortening contractions without a preceding stretch (Cavagna et al.1968; Seiberl et al. 2015), which is known as the SSC effect.

The SSC effect has been explained by mechanical and neural factors that are triggered by the stretch preceding the shortening (Seiberl et al. 2021). Mechanically, altered cross-bridge kinetics and increased passive forces contribute to the increased force capacity of muscles during active muscle stretch, which we have called transient force enhancement (tFE; Bakenecker et al. 2020). tFE could subsequently cause higher forces (Katz, 1939; Edman et al., 1978, Leonard et al., 2010) and greater elastic energy return during successive shortening. Additionally, a phenomenon known as residual force enhancement (rFE; Abbott & Aubert, 1952; Edman et al., 1982), might also enable higher forces during and following shortening (Seiberl et al., 2015; Fukutani et al., 2015a; Hahn and Riedel, 2018). As rFE is thought to be caused by increased cross-bridge forces and the engagement of the elastic element titin (Rode et al., 2009; Rassier et al., 2012; Herzog et al., 2016), these mechanisms likely contribute to the SSC effect as well (Groeber et al., 2019; Tomalka et al., 2020; Hahn et al., 2023). However, neural mechanisms could also contribute to the SSC effect. For example, stretch reflexes that are triggered during active muscle stretch, by excitatory afferent muscle spindle discharges, could facilitate and synchronize the activation of the motor units during the shortening phase of the SSC (Trimble et al. 2000). This reflex activation could thus contribute to the SSC effect by increasing muscle activation compared with pure shortening contractions (van Ingen Schenau et al. 1997; Trimble et al. 2000; Taube et al., 2012a). In contrast, the SSC effect might be attenuated by inhibitory mechanisms that are triggered during active muscle stretch (Cronin et al., 2011; Westing et al., 1991; Davey et al., 1994; Duchateau and Enoka, 2016). But as some studies did not find any neural inhibition during active muscle stretch (e.g. Hahn et al. 2012; for review see Hahn 2018), it is unclear whether there is a net inhibitory or net excitatory effect during active muscle stretch, which might originate from spinal and/or cortical sites.

To test excitability modulations at spinal and cortical sites, Gruber et al. (2009) elicited motor-evoked potentials (MEPs) and cervicomedullary motor-evoked potentials (CMEPs) in the human biceps brachii and brachioradialis muscles. These authors found smaller MEPs and CMEPs during active muscle stretch compared with fixed-end contractions, but increased MEP-to-CMEP ratios, which can be interpreted as increased cortical excitability during active muscle stretch. Further, it was shown that descending cortical drive contributes to early muscle activation during hopping SSCs, which usually was attributed to spinal stretch reflex activation (Zuur et al., 2009). Cortical and spinal excitability modulations might not just occur during active muscle stretch, but excitability modulations might persist following active muscle stretch during shortening and the isometric hold phase (i.e. steady state) following the shortening phase of SSCs. In the steady state following active muscle stretch, Hahn et al. (2012) found larger MEPs and unchanged CMEPs in the presence of rFE during maximal voluntary plantar flexion contractions, which was interpreted as increased cortical, but unchanged spinal excitability. However, Sypkes et al. (2018a) found smaller CMEPs and an unchanged MEP-to-CMEP ratio in the presence of rFE during submaximal dorsiflexion contractions, which was interpreted as unchanged cortical, but reduced spinal excitability. These two studies need to be considered with caution though, as only a relatively small number of MEPs were averaged from a relatively small sample (Frischholz et al. 2022). By averaging 20 MEPs and inhibiting the motor cortex following active muscle stretch, Frischholz et al. (2022) found that rFE was unaffected by corticospinal excitability modulations. However, this study did not assess whether corticospinal excitability was modulated during active muscle stretch or whether cortical and spinal excitability were modulated differently following stretch.

While active muscle stretch can trigger rFE, active muscle shortening can trigger residual force depression (rFD), which refers to the long-lasting reduction in muscle force following shortening (Abbott & Aubert, 1952). In the steady state following active muscle shortening, Grant et al. (2017) found unchanged MEPs and unchanged CMEPs in the presence of rFD during maximal voluntary dorsiflexion contractions, which can be interpreted as absent cortical and spinal excitability modulations. In contrast, Sypkes et al. (2018b) found unchanged MEPs, but increased CMEPs in the presence of rFD during submaximal voluntary dorsiflexion contractions, which can be interpreted as decreased cortical, but increased spinal excitability.

Based on the contradicting findings regarding potential neural modulations triggered by active muscle stretch and shortening, no clear statement can be made as to whether cortical and/or spinal excitability is altered by active muscle stretch and shortening. Further, to the best of our knowledge, no data exists on whether the combination of active muscle stretch and shortening modulates cortical and/or spinal excitability during and following SSC contractions. Therefore, the aim of this study was to investigate cortical and spinal excitability during and following the shortening phase of SSCs compared with pure shortening contractions during submaximal voluntary plantar flexion contractions. By doing this, we aimed to gain insight into possible neural mechanisms that contribute to the SSC effect. We expected an increased torque during the shortening phase of SSC compared with SHO when the soleus EMG amplitudes were matched. We also expected reduced or similar rFD during the steady state following the shortening phase of SSC compared with SHO. Finally, we expected that possible stretch-triggered changes in cortical and/or spinal excitability would be long-lasting and that these changes would be visible during the shortening and steady-state phases of SSC compared with SHO when EMG amplitudes were matched.

## Methods

### Participants

Eighteen recreational active healthy adults (seven women, eleven men, 25.9 ± 5.6 yrs, 1.79 ± 0.05 m and 75.0 ± 10.5 kg) gave written informed consent prior to participating in this study. All subjects were free of neuromuscular diseases and of musculoskeletal injuries for at least six months prior and throughout the study. In addition, participants with a known increased risk of epileptic seizures, pregnant women, participants with implanted biomedical devices and/or electronic implants (e.g. brain pacemakers, heart pacemakers) or metal particles in the skull were excluded from participation. Ethics were approved by the Human Ethics Committee of Ruhr University Bochum (EKS V 16/2021) and all procedures were in accordance with the principles of the Declaration of Helsinki.

### Torque, angle and EMG

Resultant ankle joint torque and crank arm angle were recorded from the right foot of the participants’ leg, while participants laid prone on the bench of a dynamometer (IsoMed2000, Ferstl GmbH, GER). The right foot was tightly strapped onto a footplate attachment to minimize heel lift during contractions. Hip and knee joint angles remained fully extended, and evasive movements during contractions were minimized by securing the participants’ waist with a belt. Further, particular attention was paid in monitoring the participants’ posture during the test and in avoiding head rotations; therefore, participants hyperextended their lower back and propped their head up on their hands. The lateral malleolus was aligned with the axis of rotation of the dynamometer in a neutral position (i.e., foot plate perpendicular to the shank), which we defined as 0° ankle joint angle. The range of motion was set from -10° plantar flexion (PF) to +15° dorsi flexion (DF) (Fig. 1A). Stretch velocity was set to 40°s^-1^ and shortening velocity was set to 120°s^-1^, while angular acceleration and deceleration was set to 2000°s^-2^. Resultant ankle joint torque and crank arm angles were sampled at 1000 Hz and were filtered using a dual low-pass fourth-order 20 Hz Butterworth filter and a dual low-pass fourth-order 6 Hz Butterworth filter, respectively.

**Fig. 1.**
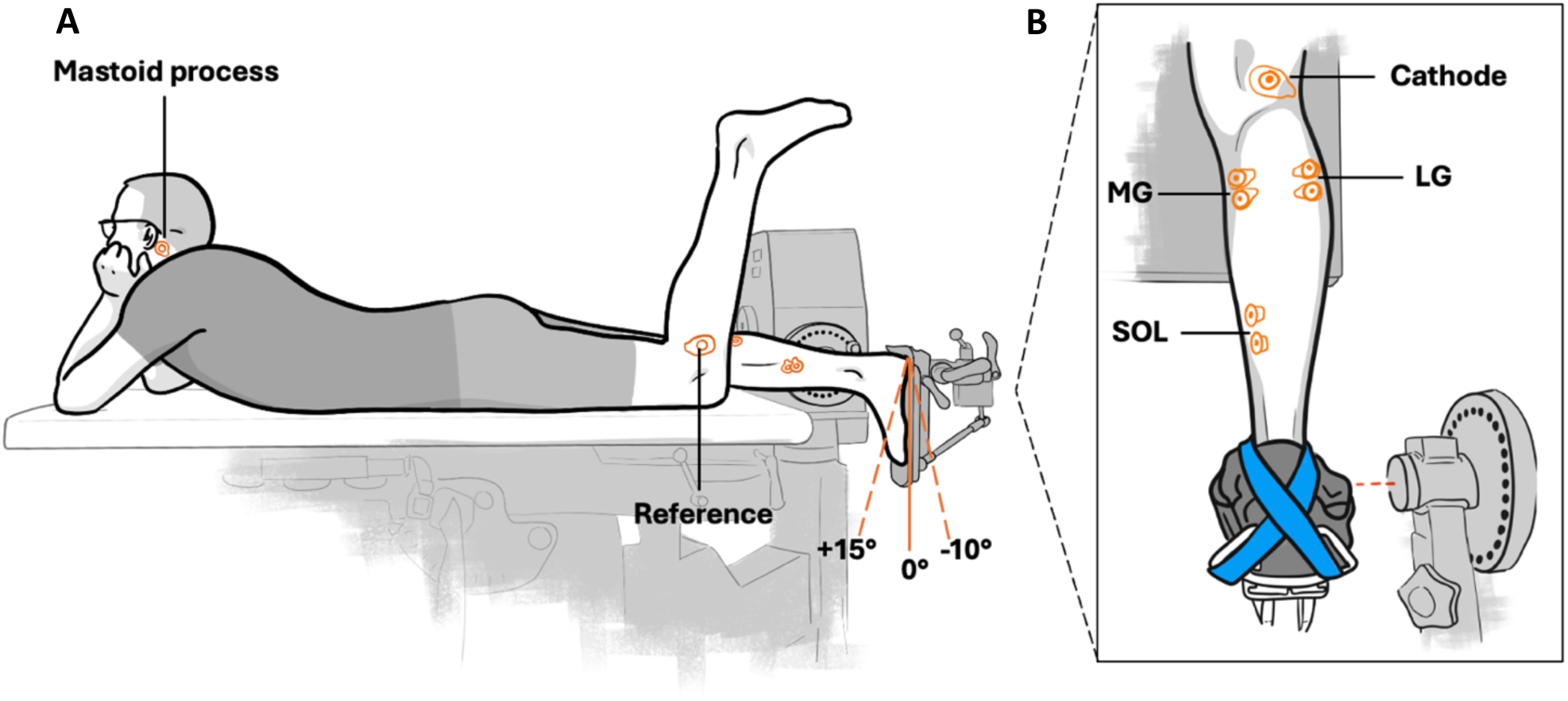
Participant set-up on the dynamometer and electrode placement on the right triceps surae. Participant laid prone on the bench of a dynamometer with the right foot strapped onto a footplate attachment. The participants’ waist was fixed with a belt (not shown). Neutral position (i.e., foot plate perpendicular to the shank) was defined as 0°. The range of motion was set from -10° plantarflexed (PF) position to +15° dorsiflexed (DF) position. Knee and hips were fully extended. Participants had to keep their head up and supported in their hands during all contractions. The reference electrode for recording muscle activity and responses to stimulations was placed on the left fibular head. Electrode for spinal cord stimulation at the cervicomedullary junction were placed over the grooves behind the mastoid processes, with the cathode on the left side of the head (**A**). Electrode placement for the recording of muscle activity and responses to stimulations of SOL, MG and LG. Electrode placement of the cathode for electric nerve stimulation within the popliteal fossa. The lateral malleolus was aligned with the axis of the dynamometer. A strap was used to firmly fix the foot to the footplate attachment (**B**).

Soleus (SOL), medial gastrocnemius (MG), lateral gastrocnemius (LG) and tibialis anterior (TA) muscle activities were recorded at 5000 Hz using surface electromyography (EMG) (Fig. 1B). EMG signals were pre-amplified 1000 times (NL844, Digitimer, UK) and high-pass filtered at 10 Hz (NL135 and NL144, Digitimer, UK) using an analog filter. EMG signals were recorded using two electrodes (8 mm recording diameter, Ag/AgCl, Covidien, Mansfield, MA, USA) placed at an interelectrode center-to-center distance of 2 cm according to international guidelines with a single ground electrode of the same type attached to the fibular head of the left leg (Fig. 1A). The skin was prepped by shaving and abrading (Nuprep, Weaver and company, Aurora, CO) prior to wiping with alcohol-based disinfectant (Sterillium, BODE Chemie GmbH, Germany) to minimize conductive resistance in the skin (Hermens et al., 1999). All data were synchronized using a 16-bit A/D card within a Power1401 data acquisition interface with Spike2 software (CED, Cambridge, UK). For visualization, SOL EMG was smoothed with moving root mean square (RMS) over a 250 ms window and normalized to 40% maximal voluntary activity (MVA) of EMG_SOL_ recorded during maximal voluntary contractions (MVC) at PF and DF.

### Stimulation techniques

Peripheral electrical nerve stimulation (ENS) was used to evoke M-waves. SOL M-waves were evoked by 1-ms single-pulse stimulations of the tibial nerve in the popliteal fossa (DS7AH, Digitimer, UK) (Fig. 1B & Fig. 2). Current passed from a cathode (Ag/AgCl electrode, Tyco Healthcare, Germany) placed within the popliteal fossa over the tibial nerve to an anode (coal rubber pad, 10.264.6 cm, Empi, USA) positioned 2 cm proximal to the patella. The stimulation site providing the greatest M-wave peak-to-peak amplitude was first located by a hand-held motor point pen (0.5 cm diameter; Compex, Germany) in PF at rest. Once the optimal stimulation site was determined, the stimulation electrode was firmly fixed to this site with tape. M-wave shape and peak-to-peak amplitude were checked for clear visibility at 0° and PF during isometric contraction at 40% EMG_SOL_. The intensity used to evoke a maximal M-wave (Mmax) of SOL during voluntary activation was multiplied by a factor of 1.2 for supramaximal motor nerve stimulation during all contractions.

**Fig. 2.**
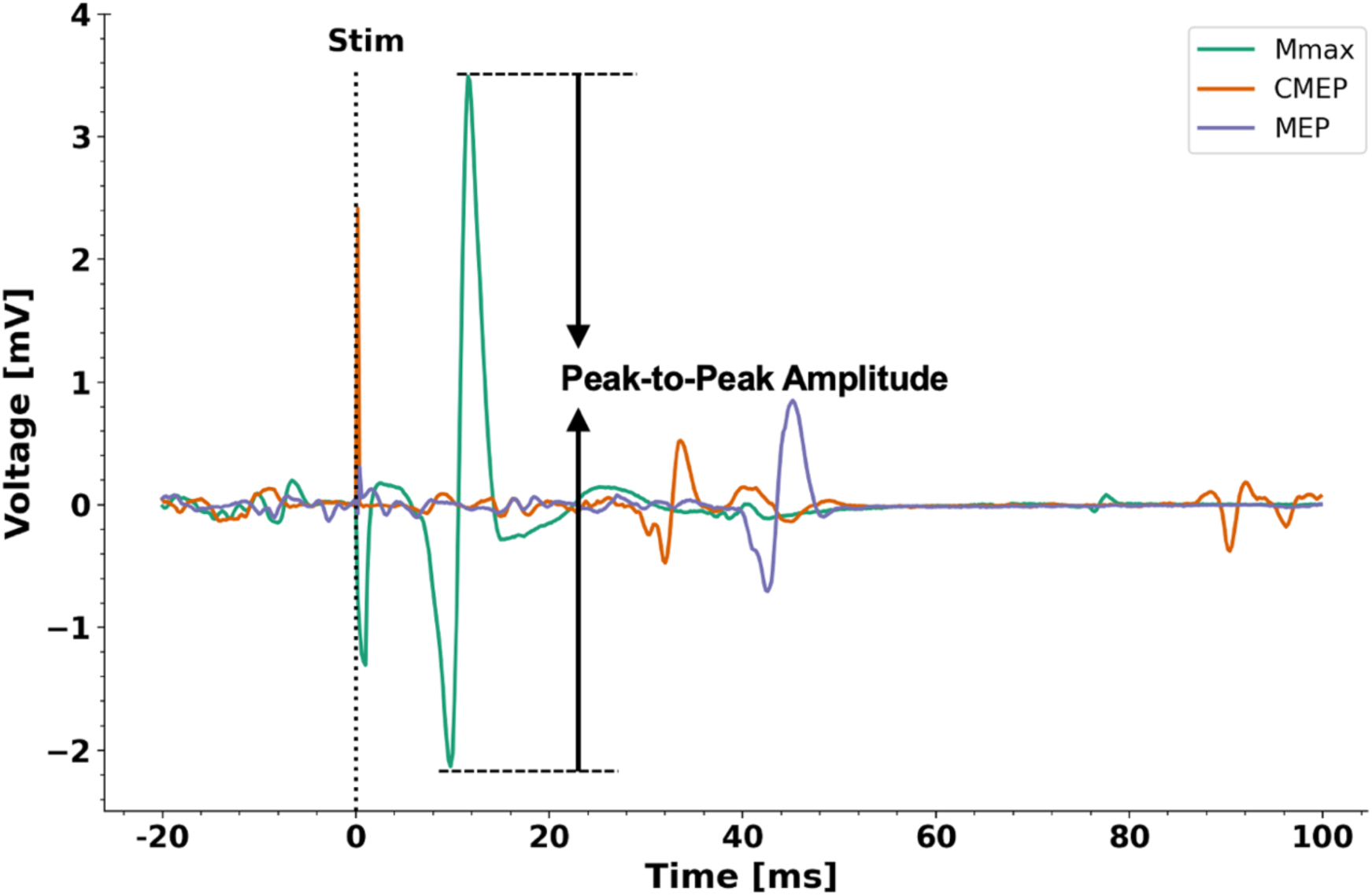
Exemplary Mmax, CMEP and MEP voltage-time responses. Individual raw EMG amplitude-time traces following stimulation from one representative participant in mV (individual muscle gains were accounted for). Mmax (green), CMEP (orange), MEP (purple). The instant of stimulation is indicated by ’Stim’. The size of the responses was calculated as the peak-to-peak amplitude.

Cervicomedullary electrical stimulation (CES) was used to evoke CMEPs in SOL (Fig. 1A & Fig. 2). CMEPs were obtained after a 100-µs single-pulse) stimulation of the spinal cord at the cervicomedullary junction using a constant voltage stimulator (D185, Digitimer, UK). Electrodes (Ag/AgCl electrode, TycoHealthcare, Germany) were placed over the grooves behind the mastoid processes, with the cathode on the left side of the head. Stimulation intensities (350-800V) were matched to 10%-30% of Mmax, while subjects maintained a submaximal contraction at 40% EMG_SOL_. CMEPs had to be clearly visible compared with the background EMG_SOL_. Due to the discomfort evoked by this kind of stimulation, stimulation intensity was not further increased than the amount each participant was able to withstand. However, if participants were unable to sustain stimulation intensities required to elicit a response equivalent to at least 10% of Mmax, they were excluded from the study.

TMS was used to evoke MEPs in SOL (Fig. 2). MEPs were elicited by a 280-µs single-pulse magnetic stimulation of the motor cortex by a double-cone coil (D-B80, MagPro Compact, MagVenture, DK), positioned to deliver anterior-posterior directed current to the brain. The coil was placed parallel to the tangent plane of the skull over the cortical motor area of the soleus, slightly left side of the vertex, which was determined through measurement of the head and then defined as the initial starting position for coil placement. Once the initial starting position was defined, several stimuli were delivered at different locations surrounding the initial starting position until the hotspot was found. The hotspot was defined as the spot where the largest MEP peak-to-peak amplitude in SOL was evoked by a single stimulation while subjects performed submaximal steady-state isometric contractions in PF at 40% EMG_SOL_. The hotspot was then marked on the skin for replication throughout the experiment. Stimulation intensity was adjusted until the peak-to-peak amplitudes of the MEPs were closely matched to those of the CMEPs.

### Protocol

Prior to testing, participants had to perform at least three familiarization sessions. During the familiarization sessions, participants had to practice all contraction conditions and were familiarized with ENS and CES. As CMEPs in lower limb muscles could not be observed in every subject (Taylor et al., 2004) non-responders (20 out of 38) were excluded during the familiarization sessions.

Participants that were included were asked to avoid any training of the lower limbs two days prior to the actual test session, as well as caffeine intake on the same day. At the beginning of the test session, participants had to perform a warm-up including ten, five, three, and two contractions at an intensity of 30, 50, 70, and 90% of perceived maximum effort, respectively. Afterwards, maximal voluntary plantar flexion torque and MVA of SOL were assessed by 2-3 MVCs in PF and DF each. Only trials with a plateau in torque and EMG_SOL_ were used for further testing, which was verified by visual inspection. Trials with the highest MVA were used to match %EMG_SOL_ based contractions.

To adjust the necessary stimulation intensities for testing, participants received visual feedback of the EMG_SOL_ on a computer monitor and were required to perform contractions at 40% EMG_SOL_ in PF in a specified order: First, Mmax was determined as a reference to adjust the sizes of the CMEPs and the MEPs to a corresponding %Mmax. Second, the simulation intensity of CES was set to obtain CMEPs, and third, based on the obtained CMEP size, the stimulation intensity of TMS was set to obtain MEPs matched to CMEP size.

Following setup of the stimulation intensities the testing started, which involved five different contraction conditions. All five contraction conditions were EMG_SOL_ controlled, which was provided as visual feedback and included the following: Two fixed-end voluntary reference contractions at PF (REF_PF_) and DF (REF_DF_) and three dynamic voluntary contractions involving pure shortening (SHO), stretch-shortening (SSC) and pure stretch (STR), which were time-matched (Fig. 3A&B). All contraction conditions (with exception of SHO) were performed with 40% EMG_SOL_ of the corresponding ankle joint angle. SSC and STR contractions started with a 2-s preactivation of 40% EMG_SOL_ before the onset of joint rotation. To match EMG_SOL_ during the shortening phase of SHO and SSC contractions, SHO contractions started with a slightly higher ankle joint-angle specific preactivation of 47% EMG_SOL_, which lasted approximately 2.5-s before joint rotation started so that shortening of SHO and SSC were time matched (see Fig. 3E). All dynamic voluntary contractions conditions (SHO, SSC, STR) were followed by an isometric hold phase at REF_DF_ or REF_PF_, respectively (Fig. 3C&D).

**Fig. 3.**
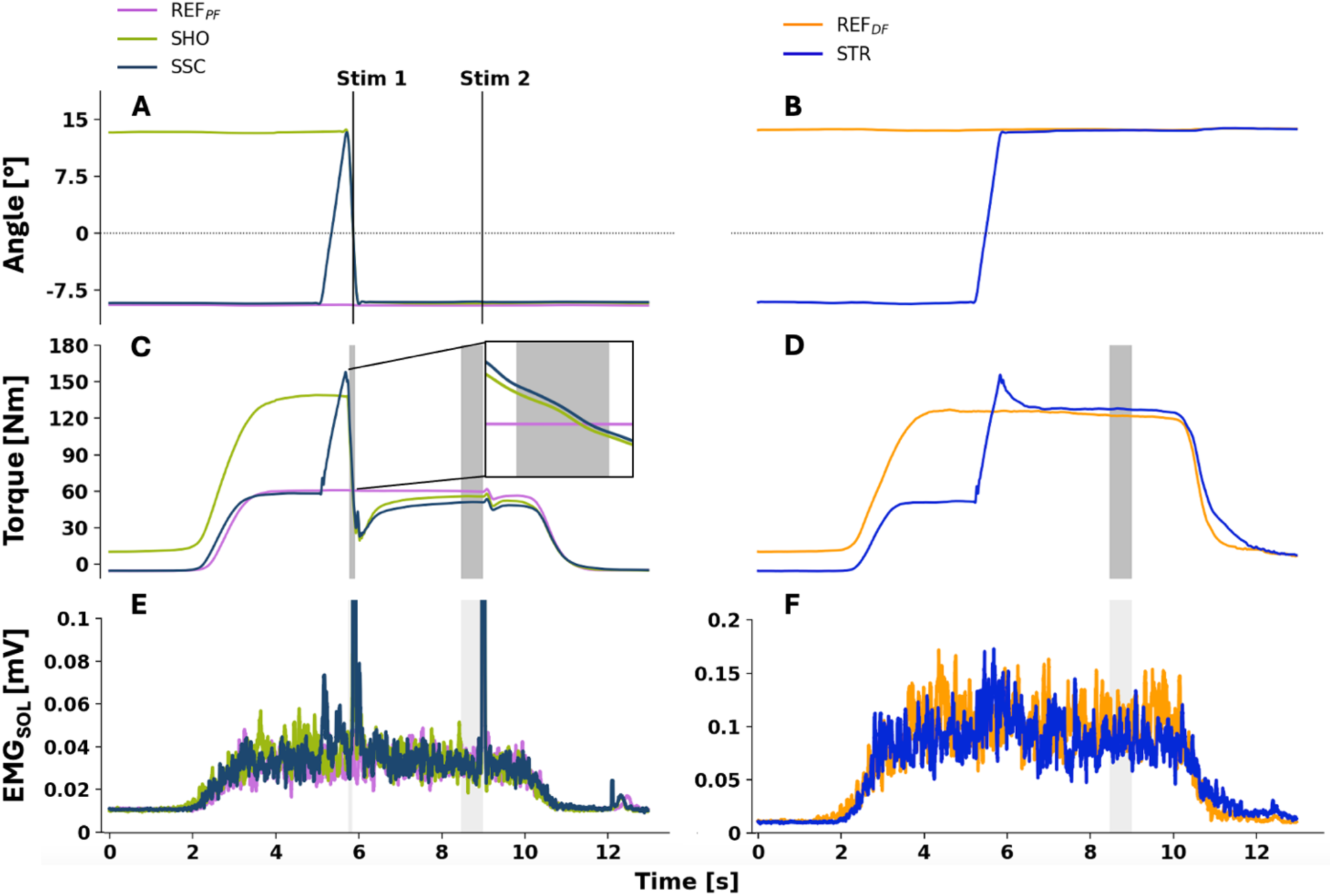
Mean crank arm angle-time (A, B), resultant torque-time (C, D) and root mean squared (200 ms) soleus muscle activity (EMG_SOL_) averaged from all participants during reference in plantarflexed position (REF_PF_), pure shortening (SHO) and stretch-shortening (SSC) (n = 15) (A, C, E) and reference in dorsal flexed position (REF_DF_) and pure stretch (STR) (n = 12) (B, D, F). The instant of stimulation during the shortening phase in SHO (green) and SSC (dark blue) is labeled as Stim 1. The instant of stimulation during the steady state of REF_PF_ (violet), SHO and SSC and is labeled as Stim 2 (**A**). No stimulation was applied during REF_DF_ (orange) and STR (blue) (**B**). The dark gray bars (**C**, **D**) indicate the time period over which torque was averaged during shortening and during the steady state for each contraction condition. The light gray bars (**E**, **F**) indicate the period over which EMG_SOL_ was averaged during shortening and during the steady state for each contraction condition and each muscle.

Further, then contraction conditions were divided into ‘stimulated’ (i.e. REF_PF,_ SHO, and SSC) and ‘non-stimulated’ (i.e. REF_DF_ and STR) conditions. During the stimulated conditions, subjects received ENS, CES and TMS superimposed on the voluntary contractions (Fig. 3C). During REF_PF_, subjects received only one stimulation while during SHO and SSC, subjects received two stimulations. Stimulation timing was set during the shortening phase of SHO and SSC, at the time when the ankle joint passed 0° ankle angle and 3 seconds after the end of the shortening phase of SHO and SSC, and at time-matched instants during REF_PF_. During STR and REF_DF_ participants received no stimulations (Fig. 3D).

All contraction conditions were randomized in blocks based on the type of stimulation (i.e. ENS, CES, and TMS) and were further divided into ’stimulated’ and ’non-stimulated’ conditions. All ’stimulated’ and all ’non-stimulated’ conditions were then further randomized based on the contraction condition. Participants had to perform three repetitions of each contraction condition for ENS, five repetitions for CES and ten repetitions for TMS for the ’stimulated’ conditions. For the ’non-stimulated’ conditions, subjects only had to perform three contractions per contraction condition. This resulted in a total of 54 submaximal contractions with superimposed stimulation and six submaximal contractions without superimposed stimulation. Participants thus received a total of 15 nerve stimulations, 25 spinal cord stimulations and 50 motor cortex stimulations. To prevent fatigue, participants had to take at least a two-minute break between contractions and longer if requested. Participants rested in PF as it was a more comfortable position.

### Data Analysis

Mean resultant torque during the shortening phase of SHO and SSC was calculated over a 105 ms time window, when shortening speed and angular displacement were constant (Fig. 3 C). The SSC effect was then calculated as the percent difference in mean torque between SHO and SSC during this time window.

During the steady state of REF_PF_, SHO and SSC, and, as well as REF_DF_ and STR, the mean resultant torque was calculated over a 500 ms time window 1 ms prior to the second stimulation instant (Fig. 3C&D). tFE was calculated as the percent difference in peak torque during the SSC stretch phase compared with the time-matched torque prior to shortening of SHO. rFD or rFE were calculated as the percent difference in mean resultant torque between REF_PF_ and SHO (rFD), between REF_PF_ and SSC (rFD), and between REF_DF_ and STR (rFE).

To determine Mmax, CMEP and MEP peak-to-peak amplitudes within each participant, a cross-correlation was performed between the averaged trials and the individual trials of each contraction condition for each muscle separately. Cross-correlations were performed over physiologically relevant time windows of 5 ms to 25 ms for Mmax, 20 ms to 50 ms for CMEPs, and 30 ms to 60 ms for MEPs (Fig. 2). Trials with a correlation coefficient >0.8 were considered as valid and included in the analysis to determine the peak-to-peak amplitude (i.e. size) of each response (Fig. 2). The mean size of each response for each contraction condition was then calculated if there were at least two (Mmax), three (CMEP), and five (MEP) valid responses.

The mean Mmax size was used to normalize the sizes of the CMEPs and MEPs during the corresponding contraction condition. The normalized CMEPs and MEPs were then averaged within each contraction condition within each participant and compared between SHO and SSC during the shortening phase, and between REF_PF_, SHO and SSC during the steady state following shortening.

EMG signals recorded from SOL, MG, LG, and TA were smoothed using a moving RMS amplitude calculation. The moving RMS window had a step size of 1 data point (i.e. 0.2 ms), a duration of 50 ms and was applied from 99 to 1 ms before the first stimulation during the shortening phase of SHO and SSC (Fig. 3E). A different moving RMS window with a step size of 1 data point and a duration of 200 ms was applied from 699 to 1 ms before the second stimulation during the steady state of REF_PF_, SHO and SSC (Fig. 3E). This same moving RMS window was applied over a time-matched period during REF_DF_ and STR (i.e. from 2300-2999 ms after the crank arm rotation; Fig. 3F).

The smoothed EMG_SOL_, EMG_MG_, EMG_LG_, and EMG_TA_ of the contraction conditions including a stimulation (REF_PF_, SHO, SSC) were normalized to Mmax of the corresponding contraction condition. The smoothed EMG_SOL_, EMG_MG_, EMG_LG_ and EMG_TA_ of the contraction conditions not including a stimulation (REF_DF_, STR) were normalized to the MVA of the respective muscle that was measured during the plantar flexion MVC in DF.

To detect potential stretch reflex activity, the raw EMG_SOL_, EMG_MG_, and EMG_LG_ data was rectified after DC removal and then averaged over all trials from either the SSC or STR conditions within each participant. In both conditions, the start of the stretch phase was defined as the onset of crank arm rotation (i.e. ankle dorsiflexion from PF to DF). The onset of crank arm rotation was defined as the last zero crossing of the crank arm position’s first derivate before crank arm velocity became and remained positive. Stretch reflexes were then visually identified for each muscle separately at different latencies from the crank arm onset. The onset of the short-latency response (SLR) was defined as the first major deflection in the mean rectified EMG of each muscle (latency ∼45ms). The end of the SLR was identified as the second mean rectified EMG amplitude peak (Ogiso et al., 2002). The subsequent EMG peaks were identified depending on their latency as mid latency response (MLR ∼70 ms), late latency response (LLR ∼90 ms), late latency response 2 (LLR_2_ ∼120 ms) (Taube et al., 2008).

### Statistical Analysis

The statistical analysis was performed with GraphPad Prism (version 10.0.3, Boston, USA), and the alpha level was set to 0.05. Results are provided as mean ± standard deviation. We used a two-way repeated-measures restricted maximum likelihood mixed-effects model to assess mean differences in normalized EMG amplitude (151 of 420 values are missing due to excluded trials) among contraction conditions and between muscles (contraction condition x muscle). Similarly, we used this model to assess mean differences in resultant torque (five missing values) among the contraction conditions and between normalized MEPs and CMEPs (contraction condition x response; 26 missing values). We used a paired t-test or a one-way repeated-measures ANOVA to assess mean differences in Mmax (zero missing values) among contraction conditions. Following a significant interaction (or a significant main effect for contraction condition when there were >2 levels), we conducted Holm-Šídák multiple comparisons to identify which conditions were significantly different. Further, a paired t-test was used to determine whether there was a significant mean difference in tFE between SHO and SSC and in plantar flexion torque during the steady state between REF_DF_ and STR.

Pearson correlation coefficients were calculated to assess the strength of the relations between the SSC effect (1) and the delta cortical excitability or the delta spinal excitability. Spinal excitability was defined as the normalized CMEP size, and cortical excitability was calculated as the difference between the normalized MEP and normalized CMEP sizes. Delta excitability was calculated as the difference in cortical or spinal excitability between SSC and SHO during shortening. Pearson correlation coefficients were also calculated to assess the strength of the relations between the SSC effect and (2) the difference in rFD between SHO and SSC during the steady state. Finally, Pearson correlation coefficients were calculated to assess the strength of the relations between the difference in rFD ((3) REF_PF_ vs. SHO, (4) REF_PF_ vs. SSC) and delta cortical excitability or delta spinal excitability.

## Results

### EMG

The mean normalized EMG_SOL_, EMG_MG_, EMG_LG_, and EMG_TA_ were not significantly affected by the contraction condition but were significantly different among muscles. However, there was no significant interaction between contraction condition and muscle. Similar results were found during the shortening phase between SSC and SHO, as well as during the steady-state phase among SSC, SHO, and REF_PF_, and STR and REF_DF_. These findings indicate successful matching of individual muscle activity levels throughout the experiment (see Table 1 for details).

**Table 1.** Normalized EMG amplitudes of the soleus (SOL), medial gastrocnemius (MG), lateral gastrocnemius (LG) and tibialis anterior (TA) muscles during the different contraction conditions. Mean ± SD values are presented in %. The mean EMG amplitude from each condition with stimulation was normalized to the maximum M-wave (Mmax) size obtained from the corresponding contraction condition from the corresponding muscle. The mean EMG amplitude from each condition without stimulation was normalized to the MVA values in the DF position of the corresponding muscle. Mean normalized EMG amplitudes were calculated during the shortening phase of SHO and SSC, as well as during the isometric steady state of REF_PF_, SHO and SSC and REF_DF_ and STR.

|  | *During shortening* |  | *During steady state* |  |  |  |  |
| --- | --- | --- | --- | --- | --- | --- | --- |
|  | *SHO* | *SSC* | *REF<sub>PF</sub>* | *SHO* | *SSC* | *REF<sub>DF</sub>* | *STR* |
| EMG<sub>SOL</sub> [%] | 1.5 % ± 0.6 % | 1.4 % ± 0.2 % | 1.1 % ± 0.4 % | 1.1 % ± 0.5 % | 1.1 % ± 0.5 % | 32.0 % ± 1.7 % | 33.3 % ± 2.3 % |
| EMG<sub>MG</sub> [%] | 2.0 % ± 0.6 % | 2.1 % ± 0.7 % | 1.7 % ± 0.7 % | 1.6 % ± 0.6 % | 1.6 % ± 0.6 % | 37.8 % ± 5.7 % | 37.6 % ± 5.9 % |
| EMG<sub>LG</sub> [%] | 2.0 % ± 0.8 % | 2.5 % ± 0.8 % | 2.0 % ± 1.0 % | 1.9 % ± 0.7 % | 2.1 % ± 1.0 % | 36.6 % ± 5.2 % | 36.4 % ± 5.2 % |
| EMG<sub>TA</sub> [%] | 0.6 % ± 0.2 % | 0.7 % ± 0.1 % | 0.6 % ± 0.1 % | 0.7 % ± 0.1 % | 0.7 % ± 0.1 % | 38.7 % ± 5.9 % | 37.4 % ± 4.5 % |

#### Torque

The mean torque during the shortening phase was significantly enhanced during SSC compared with SHO (12.0 ± 24.5%, *p* = 0.046 ; Table 2 and Fig. 4A). The mean steady-state torque was also significantly affected by contraction condition (*F*_1.22, 17.13,_ = 9.23, *p* = 0.005), but not by stimulation technique (*F*_1.52, 21.35_ = 0.40, *p* = 0.618), and there was no significant interaction between condition and technique (*F*_2.71, 37.92_ = 1.20, *p* = 0.313; Table 2 and Fig. 4B). The mean steady-state torque was significantly lower following SSC compared with REF_PF_ (-13.1 ± 3.1%, *p* = 0.006) and significantly lower compared with SHO (-7.8 ± 11.5%, *p* = 0.011). The mean steady-state torque was not significantly lower following SHO compared to REF_PF_ (-7.2 ± 8.2 %, p = 0.456). tFE was significantly enhanced during SSC compared with SHO (15.1 ± 25.2%, p = 0.003). Additionally, the mean steady-state torque was significantly enhanced during STR compared with REF_DF_ (7.3 ± 3.6%, *t* = 2.12, *p* = 0.029; Table 2 and Fig. 4C).

**Fig. 4.**
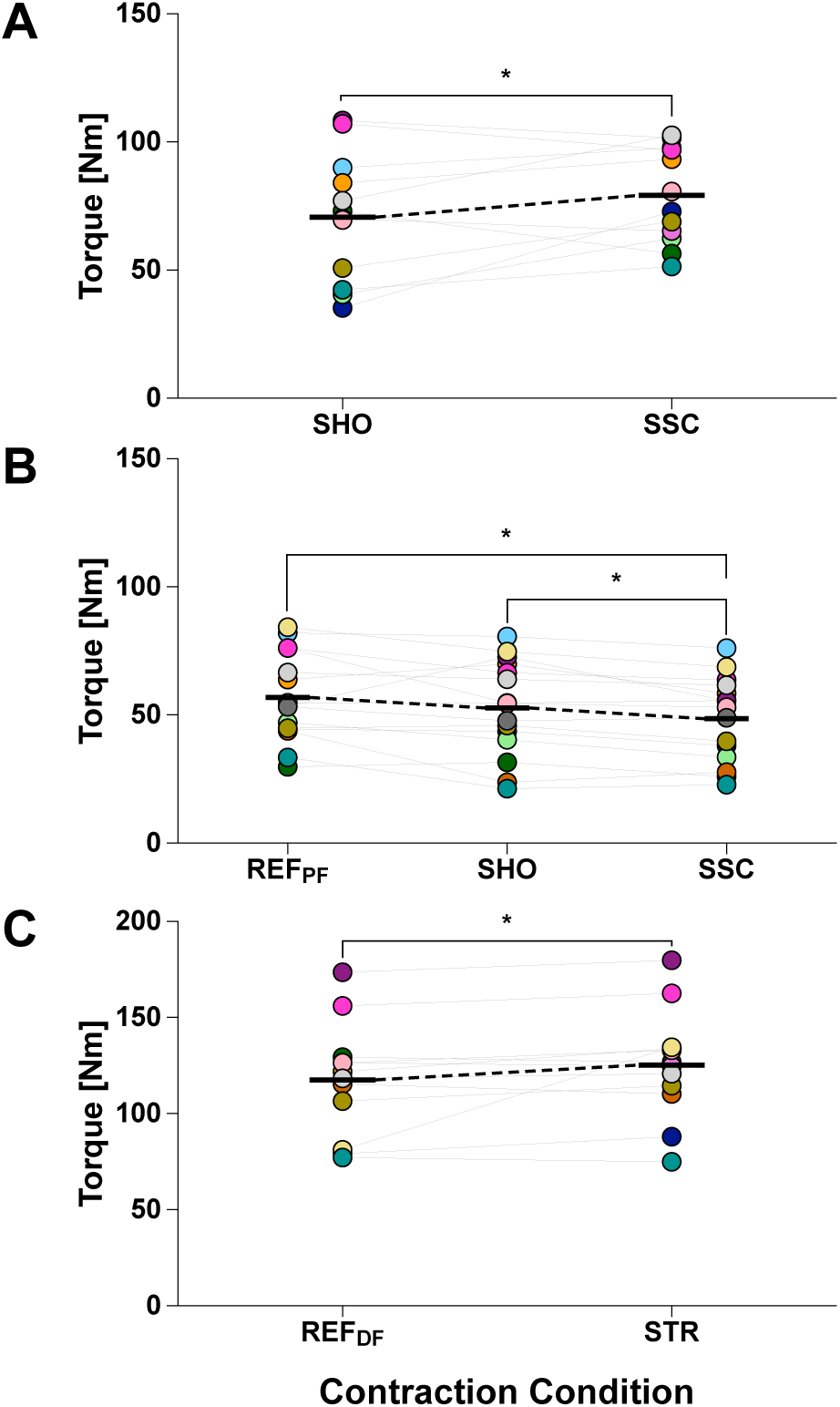
Recorded plantar flexion torques among the contraction conditions. Individual data points are presented as filled circles, mean data points as black horizontal lines connected by broken lines, and significant differences between conditions (*p* ≤ 0.05) are indicated by an asterisk. Paired values are indicated by connecting lines and identical symbol colors. During shortening, n = 12 (**A**). During the steady state, n = 15 for REF_PF_, SHO and SSC (**B**), and n = 11 for REF_DF_ and STR (**C**).

**Table 2.** Resultant plantar flexion torques. Mean ± SD values of the mean torque calculated during the shortening phase of SHO and SSC, as well as during the isometric steady state of REF_PF_, SHO and SSC and REF_DF_ and STR.

|  | *During shortening* |  | *During steady state* |  |  |  |  |
| --- | --- | --- | --- | --- | --- | --- | --- |
|  | *SHO* | *SSC* | *REF<sub>PF</sub>* | *SHO* | *SSC* | *REF<sub>DF</sub>* | *STR* |
| Torque [Nm] | 70.6 ± 24.7 | 79.1 ± 18.6* | 55.9 ± 17.0 | 53.7 ± 17.8 | 48.6 ± 16.4<sup>#</sup> | 117.6 ± 29.3 | 126.2 ± 28.0* |

\* Significantly different compared with other condition
<sup>#</sup> Significantly different compared with REF<sub>PF</sub> and SHO

### Responses

Mean SOL Mmax was not significantly different during the shortening phase between SSC and SHO (*p* = 0.454; Table 3 and Fig. 5A,C&E) or during the steady state among SSC, SHO, and REF_PF_ (*F* = 0.806, *p* = 0.442; Table 3 and Fig. 5B,D&F). Normalized SOL CMEP and MEP sizes were also not significantly different during the shortening phase between SSC and SHO (*F*_1, 6_ = 0.02, *p* = 0.885; Table 3 and Fig. 5A,C&E) or during the steady state among SSC, SHO and REF_PF_ (*F*_2, 10_ = 0.328, *p* = 0.727; Table 3 and Fig. 5B,D&F). Stretch reflexes (SLR) were only visually identified in one of fifteen participants from the SOL, but not from the MG or LG between 45 ms and 60 ms after the onset of crank arm rotation.

**Fig. 5.**
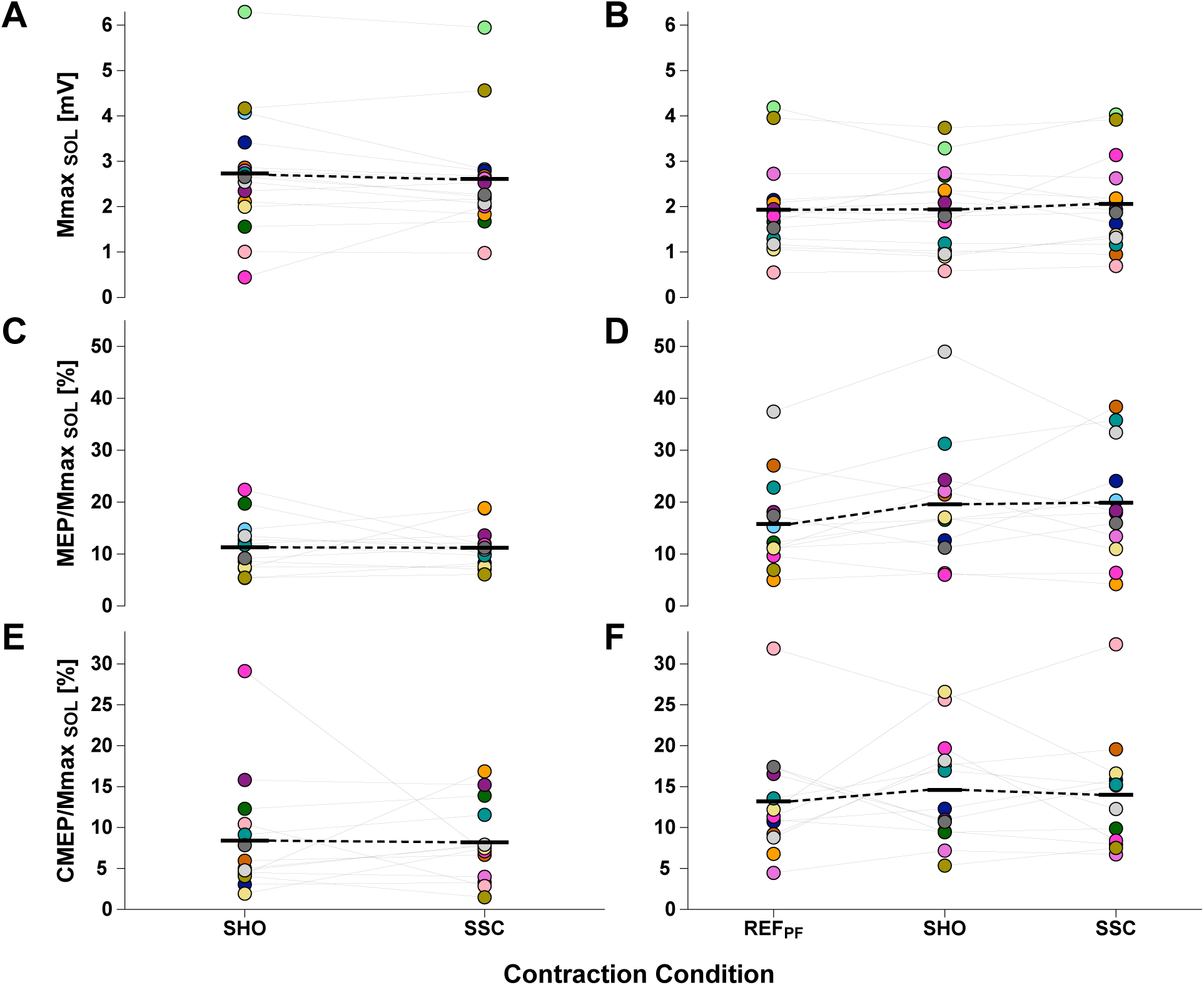
Recorded responses obtained from soleus muscle during the contraction conditions including a stimulation. Individual data points are presented as filled circles and mean data points as black horizontal lines connected by broken lines. Paired values are indicated by connecting lines and identical symbol colors. Mmax sizes are presented in mV as individual data points (n = 15) during shortening (**A**) and (n = 15) during the steady state (**B**). MEP and CMEP sizes were normalized to the Mmax size from the corresponding contraction condition. MEP/Mmax sizes are presented in % as individual data points (n = 14) during shortening (**C**) and (n = 13) during the steady state (**D**). Similarly, CMEP/Mmax sizes are presented in % as individual data points (n = 13) during shortening (**E**) and (n = 12) during the steady state (**F**).

**Table 3.**
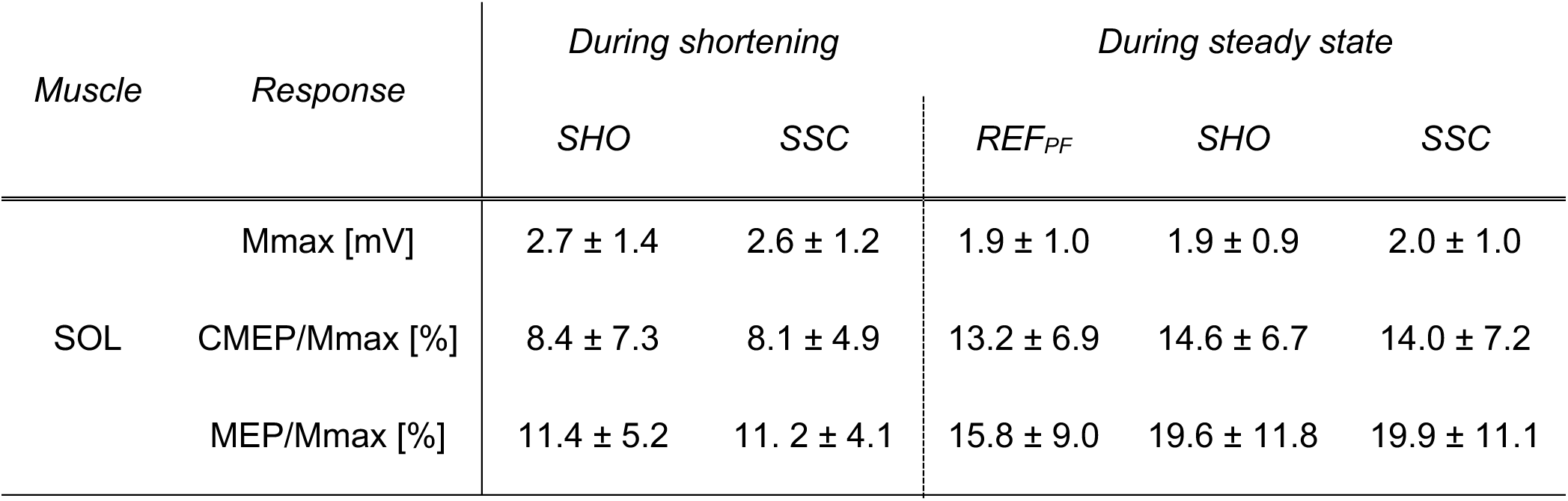
Response sizes following stimulation. Mean ± SD values of the maximum M-wave (Mmax) size, the cervicomedullary motor-evoked potential (CMEP) size, and the motor-evoked potential (MEP) size. CMEP and MEP sizes were both normalized to Mmax of the respective contraction condition. Responses were obtained during the shortening phase of SHO and SSC, and during a subsequent steady-state phase of the same conditions. Responses were also obtained during a time-matched steady-state phase during REF_PF_ at the same final ankle joint angle.

### Correlations

During the shortening phase, delta cortical excitability (CE%) and delta spinal excitability (SE%) did not significantly correlate with the SSC effect (CE%: *r* = 0.15 [95% CI: -0.52 to 0.71], *p* = 0.679; SE%: *r* = 0.16 [95% CI: -0.48 to 0.69], *p* = 0.628; Fig. 6A&B). There was also no significant correlation between the SSC effect during shortening and rFD following shortening (*r* = -0.068 [95% CI: -0.61 to 0.52], *p* = 0.833) (Fig. 7). Similarly, delta cortical excitability and delta spinal excitability did not significantly correlate with rFD for SHO (CE%: *r* = -0.52 [95% CI: -0.85 to 0.11], *p* = 0.097; SE%: *r* = 0.51 [95% CI: -0.09 to 0.83], *p* = 0.090; Fig. 6C&D) and rFD for SSC following shortening (CE%: *r* = 0.09 [95% CI: -0.57 to 0.68], *p* = 0.800; SE%: *r* = 0.23 [95% CI: -0.38 to 0.71], *p* = 0.456; Fig. 6C&D).

**Fig. 6.**
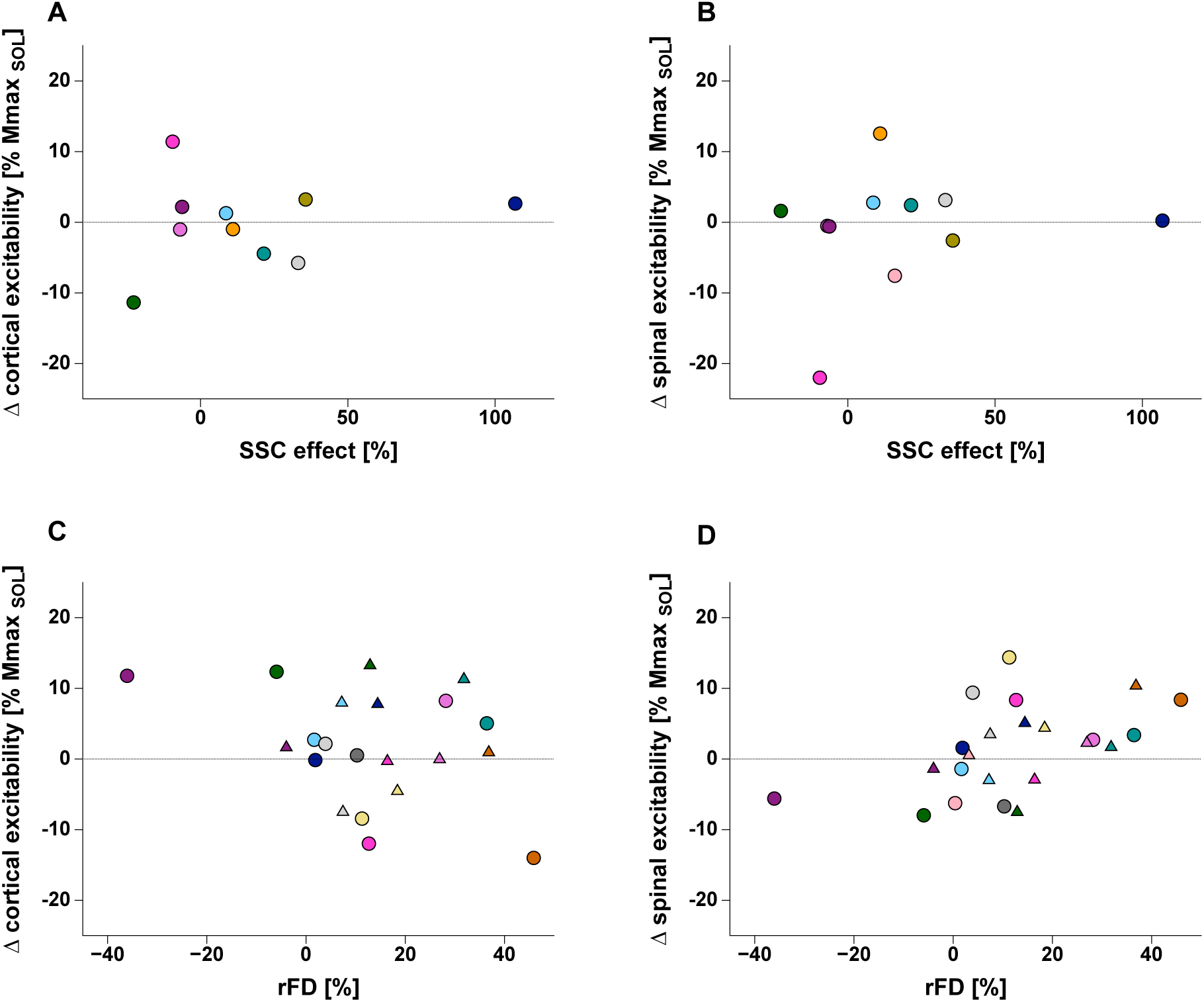
Scatter plots of delta (Δ) in cortical excitability (A, C) and Δ in spinal excitability (B, D) of the soleus muscle in relation to the SSC effect % (A, B) and in relation to rFD % (C, D). Individual data points are indicated by assigned and different colors. Delta cortical excitability calculated from the difference between normalized MEP and normalized CMEP sizes during the shortening phase (A) and during the steady state (C). Delta spinal excitability was calculated from the difference between the normalized CMEP sizes during the shortening phase (B) and during the steady state (D). Colored circles indicate the correlation between the delta excitability and SSC effect% (SSC vs. SHO) (A, B) or between rFD% (REF_PF_ vs. SSC) (C,D) and colored triangles indicate the correlation between the delta excitability and rFD% (REF_PF_ vs. SHO) (C,D).

**Fig. 7.**
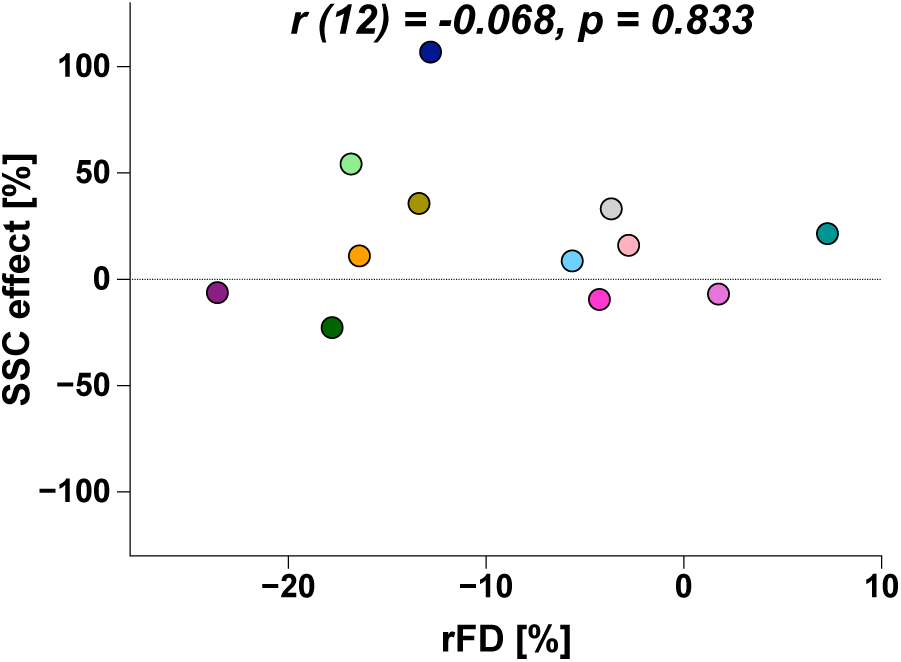
Scatterplots of the SSC effect relative to rFD. The SSC effect in % was calculated during the shortening phase of SHO and SSC and rFD in % was calculated during the steady state following shortening. Individual data points are indicated by assigned and different colors.

## Discussion

This study was the first to investigate cortical and spinal excitability during and following the shortening phase of SSCs compared with pure shortening contractions. We found a significant SSC effect, but unchanged cortical and spinal excitability during the shortening phase of SSC contractions compared with pure SHO contractions. During the steady state following shortening, we found increased rFD following SSC compared with SHO and unchanged cortical and spinal excitability. This indicates that the SSC effect is unlikely to be associated with cortical or spinal excitability modulations.

### Force enhancement during and following active muscle stretch

During and following active muscle stretch, muscle force capacity is increased compared with a fixed-end reference contraction at the same muscle length and muscle activity level. We have termed this as tFE and rFE, respectively (Bakenecker et al., 2020). To assess whether our stretch conditions elicited tFE and rFE, we performed a control condition (STR) and found significant tFE of 15 ± 25 % at the end of stretch and significant rFE of 7 ± 3%. following stretch. This latter finding aligns with previous studies showing rFE in vivo in various muscles following various stretch conditions (Lee and Herzog, 2002; Rassier and Herzog, 2005, Herzog et al. 2006, Pinniger and Cresswell, 2007; Hahn et al., 2010; Seiberl et al. 2015, Fukutani et al., 2017). Thus, our results suggest that the stretch amplitude and stretch velocity we selected were sufficient to trigger the mechanical mechanisms that result in tFE and rFE during and following active muscle stretch.

### The stretch-shortening cycle effect

We found that the mean torque during the shortening phase of SSC was significantly higher compared with SHO, and this resulted in a mean SSC effect of ∼12%, which supports our hypothesis that SSCs lead to enhanced performance during shortening. The existence of a SSC effect aligns with previous literature (Cavagna et al., 1968; and van Ingen Schenau et al. 1997), however, the size of the SSC effect in our study (12 ± 24%) was smaller. This could be because previous studies used muscle belly or peripheral nerve stimulation rather than voluntary contractions and we used a higher preload for our pure SHO versus SSC contractions. Regarding potential mechanisms behind the SSC effect, a recent study (Göcking et al. 2024) showed that the increased torque at the onset of shortening during SSCs due to active muscle stretch (i.e. tFE and its underpinning mechanisms) contributed to the observed SSC effect. This also supports the early assumption by Cavagna and coauthors (1968) that the SSC effect is partly due to contractile components, which recently was further supported by work that found a SSC effect using skinned fiber preparations (Tomalka et al. 2020, 2021).

### The history dependence of muscle action

Based on the finding that rFE was not completely abolished following shortening during SSCs, it was hypothesized that tFE and rFE share some common mechanisms and that these stretch-induced mechanisms contribute to the SSC effect (Seiberl et al. 2015). This is why we obtained steady-state torques following SHO and SSC. Although we observed rFE following active muscle stretch, we found a significant rFD of ∼13% following SSC, while the mean torque following SHO was non-significantly reduced by ∼4% compared with REF_PF_ and non-significantly larger by ∼10% than the mean torque following SSC. This is contrary to our expectations and to previous literature reporting rFE, non-significant rFD or reduced rFD following SSCs (Fortuna et al., 2017; Fortuna et al., 2018; Hahn et al., 2018). Further, our results showed no significant correlation between the SSC effect and the magnitude of rFD (Fig. 7). Taken together, these results suggest that under the given contraction conditions, the stretch-induced mechanisms underpinning rFE did not contribute much or at all to the observed SSC effect, but that the mechanisms underpinning rFD rather attenuated the SSC effect. As this is in contrast to similar previous studies using electrical stimulation, (Fukutani et al., 2015b; Fukutani et al., 2017; Hahn et al., 2018) again the type of muscle activation (i.e. electrically vs. voluntarily) might help to explain the contradicting results (Paternoster et al., 2021). Further, it was shown that the mechanisms underpinning rFE and their contribution to the SSC effect depends on the specific contraction conditions (i.e. shortening amplitude and shortening speed) (Fukutani et al., 2019; Bakenecker et al., 2022), which were not varied in this study.

### Stretch reflex during active muscle stretch

Apart from the mechanical contributions discussed above, stretch reflex activity is potential contributor to the SSC effect (Komi et al., 1997; Turner et al., 2010). However, in our study, SLR stretch reflexes were found in only one out of fifteen participants for SOL, but not for MG or LG. This is in contrast to studies reporting a significant contribution to the SSC effect by stretch reflex activity in human triceps surae muscles (Dietz et al., 1981, Zuur et al., 2009). A reason for the contrary findings might be that stretch reflex activity was usually observed during SSC-type activities such as running, hopping, or landing (Dietz et al., 1981; Komi et al., 1997; Taube et al., 2012b, Schuster et al., 2020), which involve much faster joint rotations compared with the controlled dynamometer conditions in this study. Accordingly, the specific contraction conditions might play a crucial role regarding the potential contribution of stretch-reflex activity to the SSC effect.

### Cortical and spinal excitability during muscle shortening

tFE and rFE have not only been linked to the SSC effect, but also to cortical and spinal excitability modulations (Gruber et al., 2009; Hahn et al., 2012; Sypkes et al., 2018a). This is why we investigated whether cortical and/or spinal excitability differed between muscle shortening when preceded by a fixed-end contraction during SHO and by active muscle stretch during SSC, respectively. Our results showed no significant differences in normalized MEP and CMEP amplitudes during the shortening phases of SHO and SSC, which suggests that cortical and spinal excitability remained unaltered when shortening was preceded by active muscle stretch. Further, the non-significant correlations between changes in cortical and spinal excitability and the SSC effect (Fig. 6A & 7B) indicate that cortical and/or spinal excitability modulations did not contribute to the SSC effect. Although we did not investigate potential cortical and spinal excitability modulations during the active stretch phase, our findings indicate that potential cortical and spinal excitability modulations during active muscle stretch do not persist throughout subsequent muscle shortening during SSC. This interpretation is in line with findings that corticospinal excitability remained unaltered during the steady-state hold phase in the presence of rFE following active muscle stretch (Frischholz et al. 2022). However, if cortical and spinal excitability also remained unaltered during stretch as also previously reported in literature (Hahn et al., 2012), then our findings indicate that the SSC effect is rather driven by mechanical than neural mechanisms, which is in line with previous interpretations (Fukutani et al., 2021).

### Cortical and spinal excitability following active muscle shortening

To finally investigate whether active muscle length changes led to cortical and/or spinal excitability modulations, we obtained MEPs and CMEPs during REF_PF_, and during the steady state following shortening of SHO and SSC. As the size of the normalized MEP and CMEP responses during the steady states of REF_PF_, SHO and SSC were not significantly different, our results indicate that cortical and spinal excitability remained unaltered and were not affected by any active muscle length change. Further, the non-significant correlations between changes in cortical and spinal excitability and rFD (Fig. 6C & 7D) indicate that cortical and/or spinal excitability modulations did not affect the steady-state torque following shortening. Contrary to our findings, the only other study that assessed cortical and spinal excitability in the presence of rFD (Sypkes et al. 2018b) observed a decreased cortical excitability, but an increased spinal excitability in the presence of rFD. Based on our findings, we conclude that rFD was not affected by cortical and spinal excitability modulations, but that rFD is rather due to mechanisms such as cross-bridge inhibition following active muscle shortening (Lee and Herzog, 2003; Joumaa et al., 2012; Chen et al., 2019; Hahn et al., 2023; Raiteri et al., 2024; Liu et al., 2024).

### Limitations

A few limitations need to be considered when interpreting our results. Cortical and spinal excitability modulations were only obtained at a single instance in time around the middle phase of shortening. However, cortical and spinal excitability modulations occur rapidly (Chen and Hallett 1999). Therefore, it is possible that cortical and spinal excitability modulations occurred earlier or later during shortening. Further, cortical and spinal excitability modulations as assessed by MEPs and CMEPs do not reveal potential specific interneuronal inhibitory and excitatory mechanisms (Kalmar, 2018). Also, our corticospinal excitability modulation results are specific to our setup, which involved targeting a specific hotspot with TMS using a figure-eight coil, using fixed stimulation parameters (Siebner et al., 2022), and having participants in a prone position. Finally, our findings are also specific to the controlled dynamometer conditions used, which involved stretch and shortening velocities of 40°s^-1^ and 120°s^-1^ over an amplitude of 25°, and do not preclude cortical and spinal excitability modulations during other SSC conditions.

### Conclusion

Our study provides first evidence that the SSC effect is not associated with cortical and spinal excitability modulations, but rather driven by mechanical mechanisms triggered during active stretch. Other neural mechanisms, including stretch reflex activity and preactivation did not drive the SSC effect. Future research should aim to investigate specific inhibitory and excitatory mechanisms at cortical and spinal sites to improve our understanding of motor control during SSCs.

## Data availability statement

The data from this study will be made available before publication.

## Authors contributions

L.F.R., B.J.R, T.S, W.S and D.H. contributed to the conception of the study; L.F.R contributed to acquisition and analysis of data; L.F.R., B.J.R and D.H. contributed to the design of the study, interpretation of data and drafted the manuscript; all authors contributed to revising the manuscript critically for important intellectual content and approved the final version of the manuscript. All authors agree to be held accountable for all aspects of the work, those designated as authors qualify for authorship, and those who qualify for authorship are listed.

